# Re-Evaluation of *MEFV* Gene Variants: Utility of a Gene Specific Threshold in Reducing the Number of Variants of Unknown Significance

**DOI:** 10.1101/532804

**Authors:** Matteo Accetturo, Angela Maria D’Uggento, Piero Portincasa, Alessandro Stella

## Abstract

Familial Mediterranean Fever (FMF) is an inherited autoinflammatory syndrome caused by mutations in the *MEFV* gene. *MEFV* variants are still in large part classified as variant of uncertain significance (VOUS), or with classification unresolved, posing significant challenges in the clinical diagnosis of Familial Mediterranean Fever (FMF). REVEL is a recently developed variant metapredictor tool. To reduce the number of *MEFV* variants with ambiguous classification we extracted the REVEL score for all missense variants reported at the locus specific database INFEVERS, and analyzed its correlation with expert-based classification and localization in the *MEFV*-encoded pyrin protein functional domains.

The data set of 216 missense variants was divided in four classification categories (BENIGN, VOUS, PATHOGENIC and UNRESOLVED). *MEFV* variants were plotted onto the pyrin protein, the distribution of REVEL scores in each category was computed and means, confidence intervals, and area under the receiver operating curve were calculated.

We observed a non-random distribution of pathogenic variants along the functional domains of the pyrin protein. The REVEL scores demonstrated a good correlation with the consensus classification of the International Study Group for Systemic Autoinflammatory Diseases (INSAID). Sensitivity, specificity, and accuracy were calculated for different cutoff values of REVEL scores and a gene-specific threshold was computed with confidence boundary limits. A REVEL score of 0.298 was the best performing cut-off to reclassify 96 *MEFV* gene variants previously of uncertain significance or unsolved thus reducing their proportion from 61.6% to 17.6%.

In conclusion, the combination of available expert information with highly sensitive predictor tools yields to more accurate interpretation of clinical consequences of *MEFV* gene variants. This approach should bring to a better genetic counseling and patient management.

**Author summary:** We aimed to refine *MEFV* gene variants classification using the metapredictor REVEL. We demonstrate that a gene-specific threshold is effective for accurate variants’ classification. Using this threshold, we reduced significantly the proportion of *MEFV* variants with an ambiguous classification. The proposed classification could represent a useful resource for variant interpretation in the context of FMF diagnosis.

## Introduction

Familial Mediterranean Fever (FMF) is the most common of the Hereditary recurrent fevers (HRFs) with an estimated prevalence of 1 in 1-5/ 10 000. FMF has been first described more than sixty years ago [1,2], and since the identification of the causative gene (*MEFV*, NM_000243.2) it has been intensively studied [3,4]. The *MEFV* gene encodes for an 86 kDa protein (Pyrin or marenostrin, TRIM20) prevalently expressed in immune cells (neutrophils, eosinophils, monocytes, dendritic cells and synovial fibroblasts), and is related to abnormal response of the innate immune system and IL-1β during periodic attacks of fever and serositis [5,6]. In spite of several reports describing large FMF patients’ datasets, a complete understanding of the pathogenic mechanisms responsible for this disease has not been reached yet. Recent studies suggested the possibility of dominant inheritance, at least for some mutations. This hypothesis is casting doubts on the traditional autosomal recessive mode of inheritance [7–10], a paradigm in FMF for long time. For FMF, a well-curated locus specific database is available at INFEVERS (https://infevers.umai-montpellier.fr/web/), containing genotype and clinical information for more than 500 variants. Recently, the International Study Group for Systemic Autoinflammatory Diseases (INSAID) has critically reviewed the clinical information for 858 variants in the four genes (*MEFV*, *TNFRSF1A*, *NLRP3* and *MVK*) responsible for hereditary recurrent fevers (HRFs) reaching a consensus classification for 94% of variants analyzed [11]. The notable exception was represented by the *MEFV* gene, where only 55.7% of the variants could be successfully classified with almost a third of them being variants of unknown significance (VOUS). As a result, clinicians face the harsh reality of frequently receiving an inconclusive genetic referral for FMF diagnosis. Internationally recognized guidelines have been reported by the American College of Medical Genetics (ACMG) and the Association for Molecular Pathology (AMP) [12], providing a five-tier classification of genetic variants in: pathogenic (P), likely pathogenic (LP), variant of uncertain significance (VOUS or VUS), likely benign (LB), and benign (B). However, since even the mode of inheritance in FMF families has been recently jeopardized by several reports of apparently dominant mutations, the ACMG/AMP guidelines might be of limited value in FMF. In this work, we used REVEL (a recently developed ensemble method which has been demonstrated to outperform most commonly used web-based predictors) [13], to improve the classification of many unsolved, unclassified, or of uncertain significance *MEFV* gene variants. We also report a non-random distribution of variants along the coding sequence of the *MEFV* gene. This observation possibly identifies novel hotspots for variants with a putative pathogenic role.

## Results

### Non-random distribution of variants

The REVEL score was extracted for all the 216 missense variants present at the INFEVERS database. Each variant was mapped along the coding sequence of *MEFV* gene, and with respect to the functional domains of the pyrin protein (Fig 1). We observed that PATHOGENIC variants (as defined in the methods) clustered in the SPRY and zf-B_box domains. In contrast, BENIGN variants were markedly underrepresented in functional domains of the pyrin protein being localized, in large majority, in the exon 2 of the *MEFV* gene. Finally, VOUS occurred along the entire coding sequence of the MEFV, whereas distribution of UNRESOLVED variants appeared to mostly overlap with the PATHOGENIC one.

**Fig 1.**
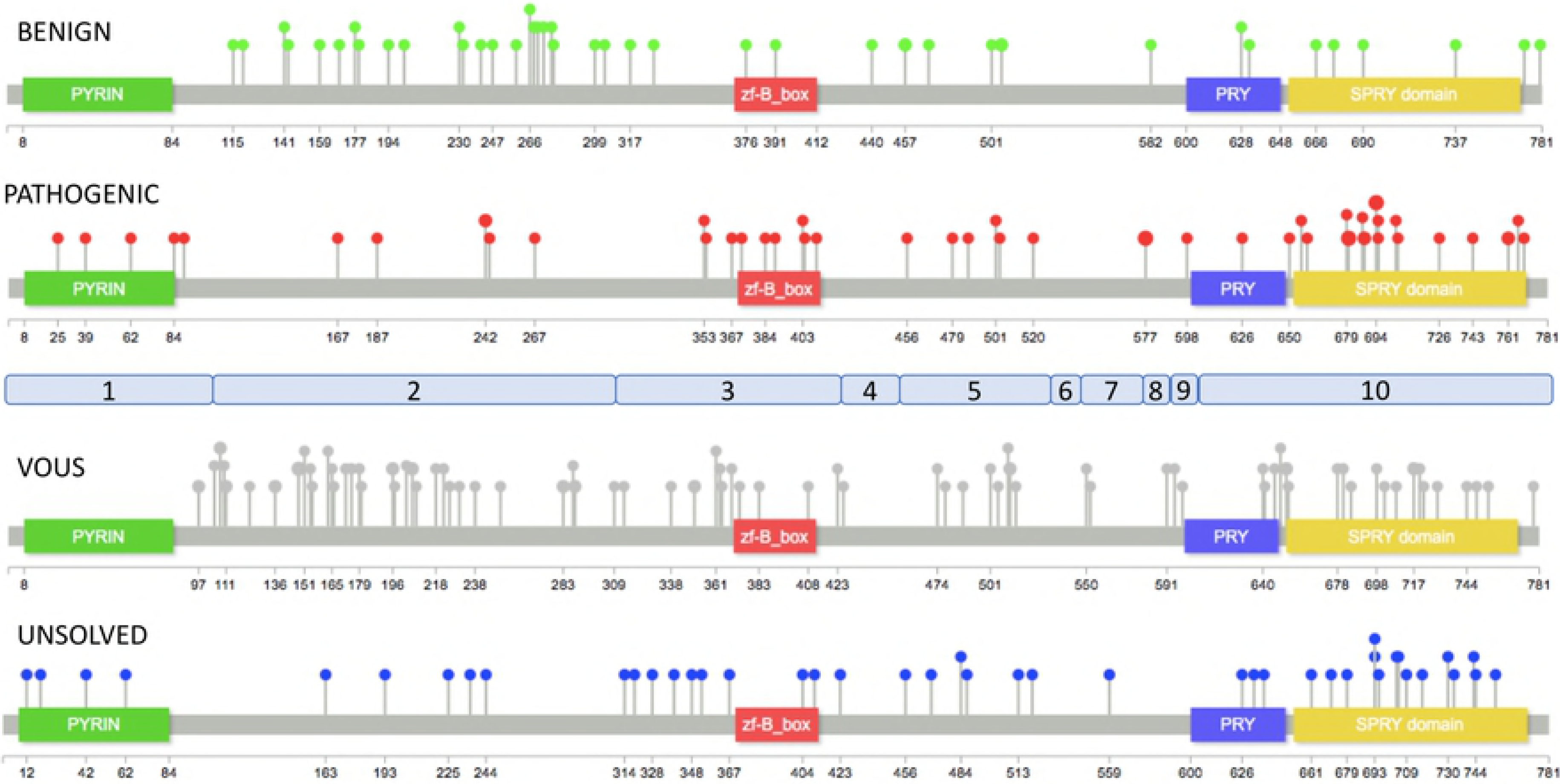
Distribution of INFEVERS variants along Pyrin Protein. Distribution of 216 MEFV INFEVERS missense variants onto the Pyrin protein. Pops size is proportional to the number of variants at a single aminoacid. In light blue are shown the exons of the *MEFV* gene.

### REVEL variants score

The means of REVEL scores for BENIGN (n=41, 0.18), VOUS (n=89, 0.25), PATHOGENIC (n=41, 0.39) and UNRESOLVED (n=44, 0.31) variants showed statistically significant differences among the four categories (F 21.04 p<0.0001, ordinary one-way ANOVA) demonstrating an appreciable level of correspondence between the INSAID consensus-based variant classification and an in silico computational scoring method such as REVEL (Fig 2). The results of unpaired t-tests among each category demonstrated statistically significant differences between BENIGN and UNRESOLVED, BENIGN and PATHOGENIC, and VOUS vs PATHOGENIC (Fig 2). Only one variant classified by INSAID as benign or likely benign had an unexpectedly high pathogenic score (E230Q, 0.737, identified as an outlier using ROUT method). In contrast, 28 of 41 INSAID pathogenic or likely pathogenic variants had a score below the REVEL threshold for pathogenicity (<0.4).[15] Worthy of note, the M694V variant, considered to be responsible for the most severe FMF phenotype, barely reached the threshold for pathogenicity (REVEL score 0.403). Finally, the area under the ROC curve (AUC) was 0.879 (*p*<0.0001, Fig 3)

**Fig 2.**
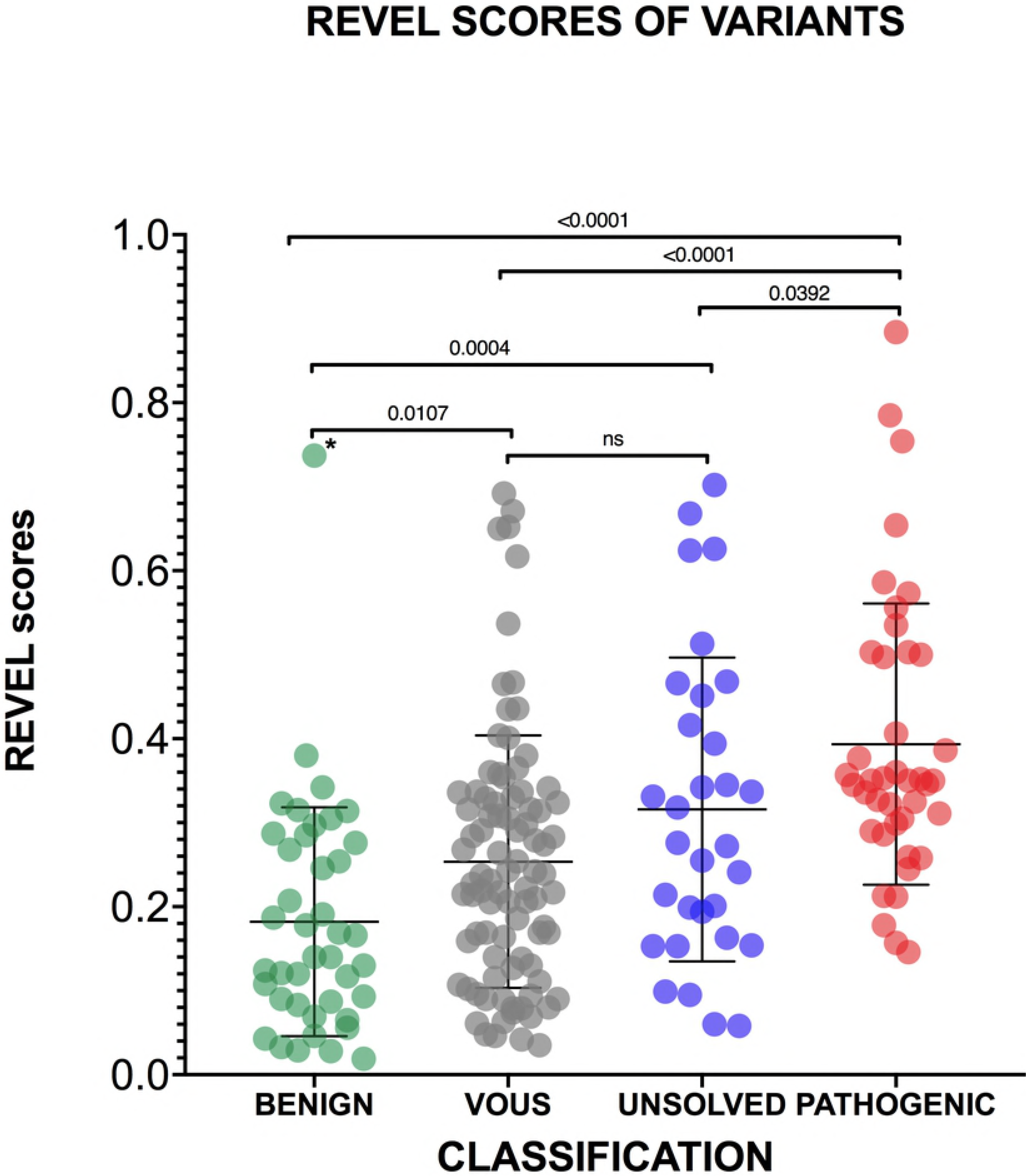
REVEL scores of MEFV variants. The mean and standard deviations are reported for each category. Unpaired *t*-tests were performed among all categories, two-tailed P values after Tukey’s post hoc analysis for multiple comparison are shown. An asterisk denotes the only variant identified as outlier (E230Q).

**Fig 3.**
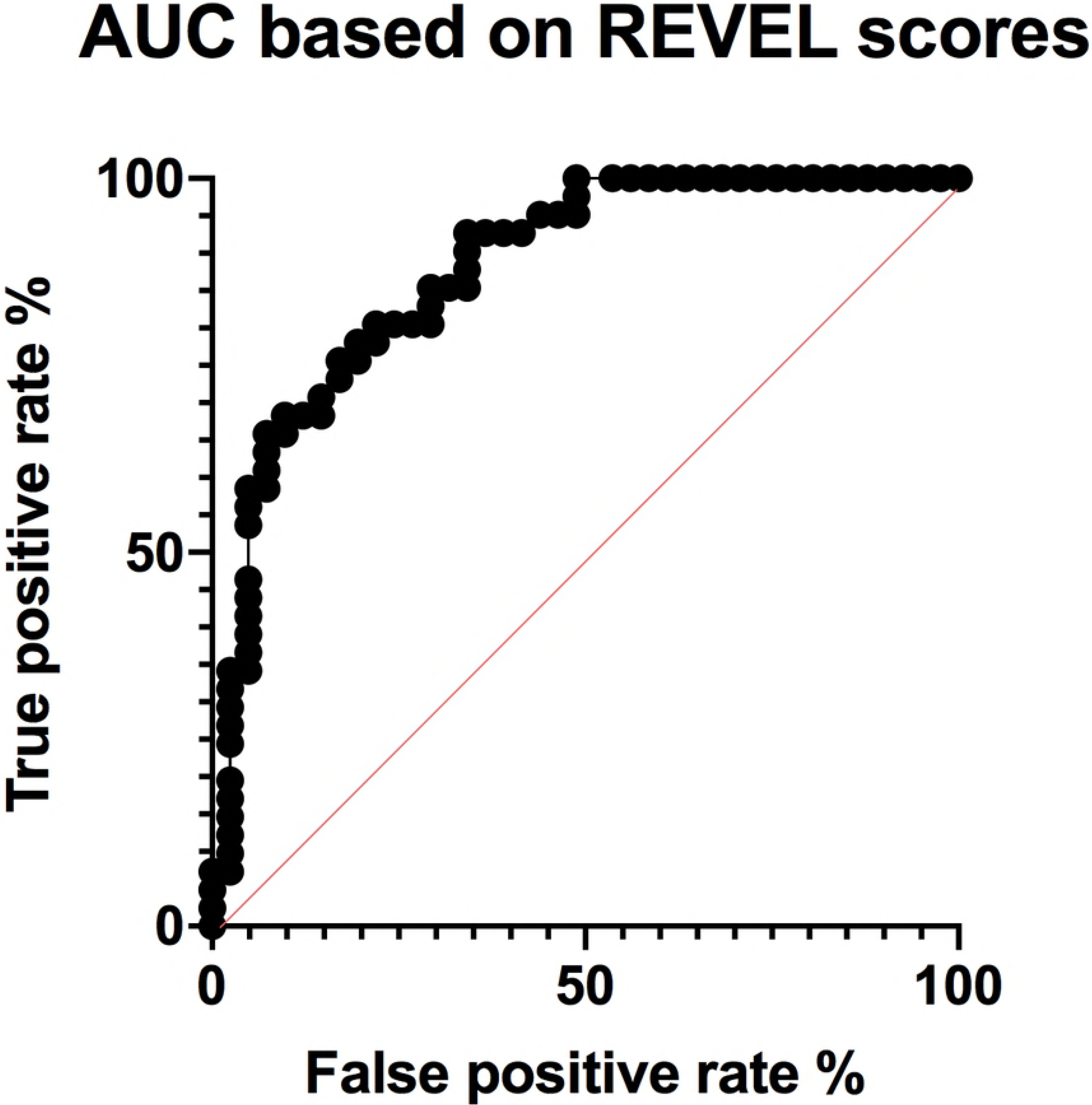
Area under the ROC curve for REVEL scores. The area under the ROC curve (AUC) is shown based on REVEL scores for the 41 BENIGN and 41 PATHOGENIC MEFV gene variants corresponding to the INFEVERS classes benign and likely benign, pathogenic and likely pathogenic, respectively.

### Development of a new classifier and reclassification of MEFV variants

Since the REVEL precision in scoring variants was markedly different for BENIGN and PATHOGENIC categories we hypothesized that *MEFV* gene variants with putative pathogenic effects may act more subtly than variants used to train REVEL. Therefore, we decided to build a model trained on a dataset formed exclusively by the MEFV variants classified by INSAID panel as BENIGN or PATHOGENIC.

Of the five algorithms tested LDA had the best performance, and was therefore further explored on the validation set. Prediction results on this set demonstrated an accuracy of 75%, with a specificity of 87.5%, and a sensitivity of 62.5%, and identified an approximate cut-off value of 0.29.

Next, we focused on the unclassified variants and aimed to assess their pathogenicity based on this previously trained model. To further optimize the performance of this new cut-off, we calculated the sensitivity and the specificity for different REVEL score thresholds (Fig 4).

**Fig 4.**
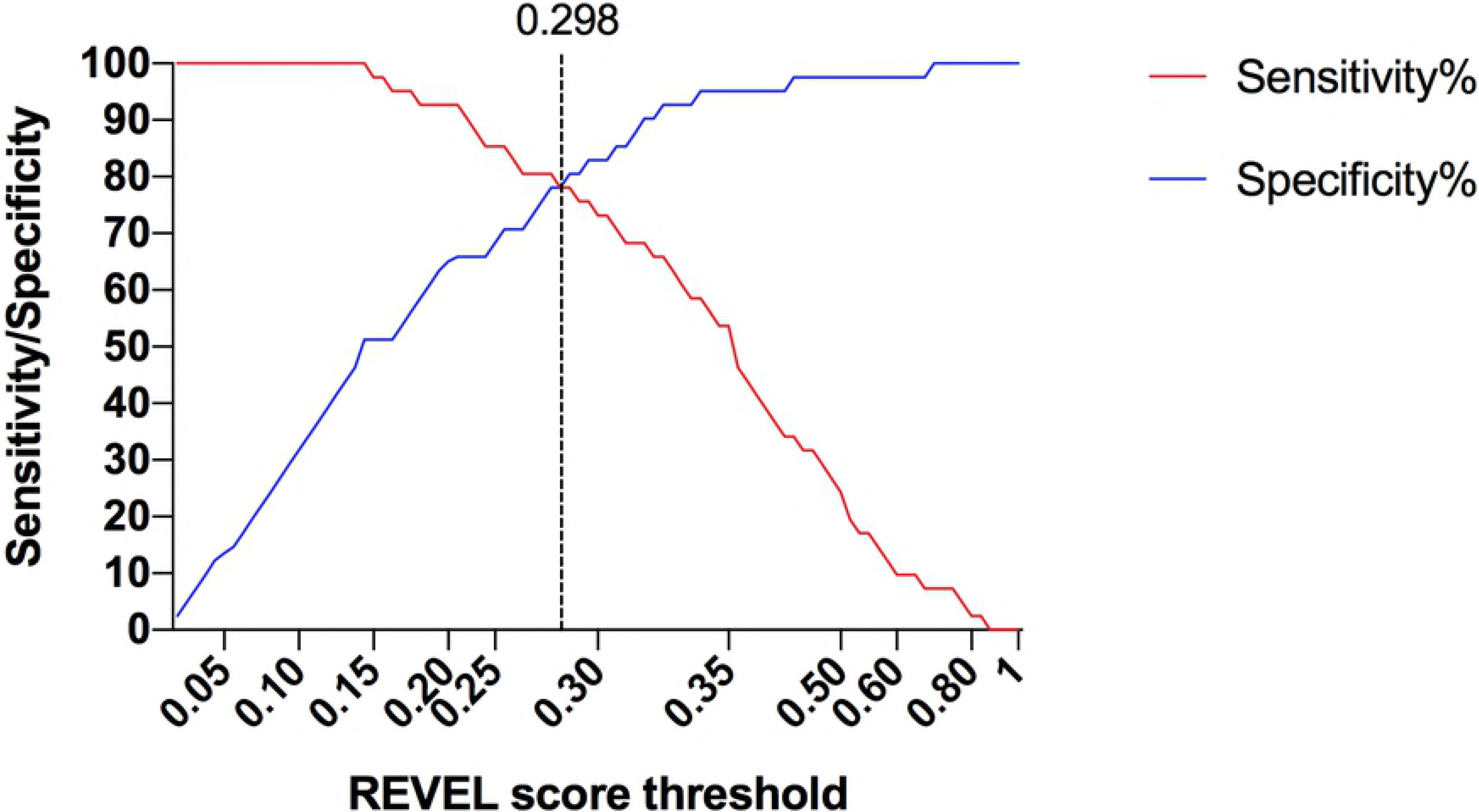
Sensitivity and specificity of different REVEL scores thresholds. Different REVEL score values were used as thresholds to assess 41 BENIGN and 41 PATHOGENIC variants of the Pyrin protein. The dotted line indicates the best performing REVEL threshold.

A nearly identical REVEL score of 0.298 yielded the best performance as cutoff, showing the highest accuracy (80.2%), positive predictive value (PPV, 83.8%) Matthews’ correlation coefficient (60.8%) and Youden index (58.2%) values, with a sensitivity of 76% and a specificity of 83%, estimated on the entire dataset of BENIGN and PATHOGENIC variants (S1 file). Thus, when using 0.298 as the cutoff value, 34 of 41 BENIGN variants had a score under the threshold for pathogenicity, while 31 of 41 PATHOGENIC variants had a score above the cutoff. The classifier we developed had an overall AUC of 0.878 (*p*<0.0001).

Then, we re-classified all the 89 “variant of uncertain significance”, 31 “unsolved” and 13 “not classified” variants from the INFEVER database with more precision, using the information deriving from the descriptive statistics of the BENIGN and PATHOGENIC REVEL scores variants to narrow the VOUS to a smaller fraction. To move from binary to graded classifications, all variants with a REVEL score lower than 0.225 (upper 95% confidence interval of the BENIGN mean) were classified as BENIGN, while variants with a score higher than 0.340 (lower 95% confidence interval of the PATHOGENIC mean) were classified as PATHOGENIC. The remaining variants (0.225≤ REVEL score ≤0.340) were considered VOUS.

Applying these criteria to the 133 VOUS and UNRESOLVED variants, 61 and 35 could be reclassified as BENIGN and PATHOGENIC, respectively (S1 Table and Fig 5). The same criteria changed the classification of 3 likely benign variants into PATHOGENIC (including the E230Q variant) and 5 likely pathogenic into BENIGN (S1 table). Notably, all but one of these 8 variants were only provisionally classified by INSAID experts.[11]

**Fig 5.**
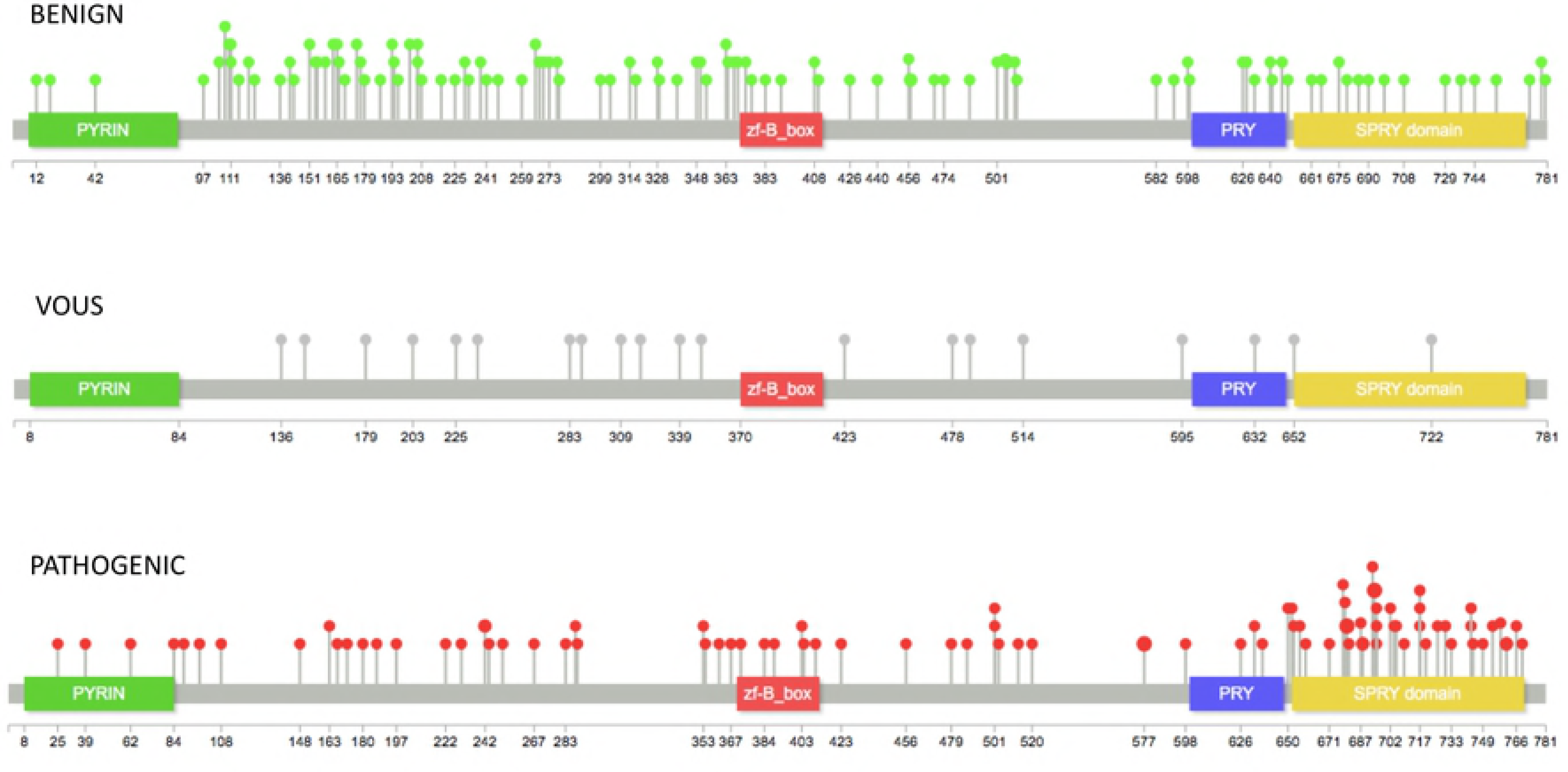
Distribution of MEFV gene variants after reclassification. Distribution of 216 MEFV missense variants onto the Pyrin protein after reclassifying VOUS and UNSOLVED variants. Pops size is proportional to number of variants at single aminoacid positions.

Finally, we mapped the newly classified variants along the MEFV protein amminoacid sequences which confirmed the preferential distribution of pathogenic variants in the SPRY and zf-B_box domains (Fig 5).

## Discussion

The advent of massive parallel next generation sequencing technology has greatly increased the power and the sensitivity of genetic testing in the clinical setting. However, the vast amount of genomic data generated poses the significant challenge of accurately assessing the pathogenicity of identified variants. While many bioinformatic resources are currently available to aid sequence variant interpretation, several inconsistencies are present in variant classification across laboratories and reference databases [16,17]. The ultimate diagnosis of FMF is linked to life-time therapy in order to prevent the complications due to secondary amyloidosis, and to definitively improve patients’ quality of life [18]. Thus, FMF presents all the above-mentioned challenges in addition to still developing effective clinical criteria for identifying truly affected patients [19,20].

In fact, of the 119 *MEFV* missense variants present in ClinVar only 24 have an ACMG classification different from VOUS or conflicting interpretations. Therefore, geneticists dealing with post-test counselling in FMF, find little help in the largest available database of clinical variation in human genome. Recently, an expert consensus panel classified the clinical significance of gene variants identified in four genes causing hereditary recurrent fevers (*MEFV*, *TNFRSFS1A*, *NLRP3* and *MVK*) [11]. However, 30% of MEFV gene variants remained unclassified or with uncertain significance.

With this initial limitation in mind, we designed the present analysis. First, by mapping variants along the coding sequence of the *MEFV*, we confirmed that pathogenic changes cluster in the SPRY domain. Two further domains, the PYRIN domain (also denoted as PYD) and the zf-B_box domain, appear to host an excess of likely pathogenic variants, although differences did not reach statistically significant difference.

The N-terminal PYRIN domain is present in more than 20 proteins modulating apoptosis and inflammation and is responsible for interaction, among others PYRIN domain-containing proteins, with the apoptosis-associated speck-like protein with a CARD (ASC) [21,22]. The zf-B_box domain is necessary and sufficient for binding the proline serine threonine phosphatase-interacting protein [PSTPIP1, or CD2-binding protein 1 (CD2BP1)]. This protein is involved in cytoskeletal organization, and its mutations cause pyogenic arthritis, pyoderma gangrenosum, and acne (PAPA), a different autoinflammatory disorder [23]. In contrast, benign variants occur largely in non-functional regions of the pyrin protein.

Next, we used REVEL, which has been demonstrated to outperform most of the available predictors, to score both classified and unclassified variants of the *MEFV* gene. When using the originally reported best threshold [13], REVEL scores were almost perfect in identifying the benign and likely benign variants (97.2%) while only 13 pathogenic/likely pathogenic variants (31.7%) demonstrated a score above the threshold for pathogenicity. We hypothesized that mechanisms conferring pathogenic effects to *MEFV* variants are subtler than mechanisms of disease-causing variants used to train many prediction algorithms including REVEL.

Indeed, the mechanisms by which *MEFV* variants cause symptoms onset are poorly understood. In a mouse model, human *MEFV* knock-in variants act as gain-of-function mutations although severity varied inversely with respect to human phenotypes [24]. More recently, in FMF patients a small set of MEFV pathogenic variants has been shown to act as hypermorphic mutations with a gene dosage effect [25]. Surprisingly, in-vitro experiments have demonstrated that FMF mutations (including M694V) render pyrin inflammasome insensitive to colchicine, while this drug is the mainstay treatment in FMF patients [26]. Although many variants with a putatively pathogenic role lie in the C-terminus of the pyrin protein, no clear genotype-phenotype correlations have emerged so far.

A further proof of the complex mechanisms leading to pathogenicity in FMF is demonstrated by the markedly different REVEL scores variants interesting the same aminoacid. For instance, the well-known polymorphism E148Q has a REVEL score of 0.274 while for the variant E148V the computed score of 0.617 has a clearly pathogenic value. Similar differences recur at five different positions along the *MEFV* coding sequence. Notably, the four most common MEFV variants on FMF chromosomes (M680I, M694V, M694I and V726A) would be all classified as neutral should the most stringent REVEL threshold score being used [27,28].

Therefore, using a classifier trained exclusively on *MEFV* variants unambiguously classified by INSAID as PATHOGENIC or BENIGN, we computed a novel REVEL threshold for pathogenicity which allowed us to propose a novel classification for all the *MEFV* variants previously considered as VOUS or UNSOLVED. Indeed, a larger dataset of variants with a definitive classification as pathogenic or benign would contribute to developing more accurate statistical models to predict the pathogenicity of variants of uncertain significance or newly identified.

The development of reliable functional assays able to assess precisely the effects *MEFV* gene variants on the pyrin inflammasome should help in expanding the set of variants with a robust classification.

The ACMG/AMP 2015 guidelines recommends that the usage of in silico predictors should have only a supporting role in establishing the pathogenic role of variants identified during genetic testing. In addition, these widely followed guidelines suggest that concordance among multiple algorithm-based predictions should be sought. However, while several metapredictors incorporating many of these older algorithms have been demonstrated to perform robustly in large heterogenous datasets, first generation algorithms such as SIFT, Polyphen, Mutation Taster, CADD and Provean are prevalently used even in recent work [29–31].

A further consideration should be mentioned about the thresholds used from the *in-silico* tools to predict functional consequences. Most of these predictors returns both a score and a prediction for the submitted variant. However, many others simply return a numeric value without any suggestion on the forecasted impact. Therefore, is up to the submitters to frame the returned score in a putative pathogenic classification and translate it in clinical recommendations for specialists and patients. REVEL, belongs to the second class of predictors, and two different threshold values for pathogenicity have been proposed by authors using this metapredictor in their studies while a threshold was not given in the original paper [13,15,28].

In this work, we identified a *MEFV* gene-specific threshold since causative mutations in FMF appears to act through a mechanism that can’t be considered neither loss-of-function (LOF) and gain-of-function (GOF). Variants leading to either LOF or GOF represent the large majority in the training datasets used to develop prediction algorithms. In contrast, *MEFV* mutations, according to a recent study seem to be hypermorphic mutations conferring to the mutant pyrin protein an enhanced reactivity to yet unknown triggers of the inflammasome [25]. Therefore, variant interpretation in FMF appears to be the prototypical case for a gene-specific calibration of prediction algorithms, as suggested by recent work [28].

In addition, based on recent analyses of mutations present in locus-specific databases (LSDB), we found that erroneous interpretations of the mechanisms responsible for the causative role of several variants can lead to misuse of prediction tools [32]. This appears to be the case also for FMF, where nine different frameshifting or nonsense variants are currently classified at INFEVERS as pathogenic or likely pathogenic in apparent contrast to the described hypermorphic nature of *MEFV* mutations [25]. Thus, critical assessment of genetic information is fundamental for translating precision medicine in everyday practice.

A limitation of our study is the small number of variants with a validated classification used to train our classifier. However, this constraint could be only overcome with an expanding set of variants identified in patients with a firm clinical evaluation that would certainly help in further narrowing the number of VOUS variants in the *MEFV* gene.

Most in silico predictors have been trained on thousands of gene variants, possibly underscoring the diverse molecular mechanisms responsible for pathogenicity in many genes. Thus, using a locus-specific scoring system may result in improved specificity and sensitivity at least for some disease-causing genes.

In summary, combining validated clinical information with the development of a gene-specific variant classifier we aimed to provide an improved framework for variant interpretation that should help in reducing the uncertainty in FMF diagnostic and would benefit genetic counsellors for a better patient management.

## Methods

### Dataset of missense variants

The dataset is based on all the 216 missense variants reported in the INFEVERS database (https://infevers.umai-montpellier.fr/web/) last accessed on 30 December 2018. We excluded from the following statistical analyses all the synonymous changes, terminating and noncoding variants, insertions, duplications and indels. We merged the ‘pathogenic’ and ‘likely pathogenic’, ‘benign’ and ‘likely benign’, “unsolved” and “not classified” variants (as classified at the INFEVERS locus specific database) in the ‘PATHOGENIC’, ‘BENIGN’, and ‘UNRESOLVED’ categories, respectively. Notably, of the 216 missense variants listed at INFEVERS only one “benign” and six “pathogenic” variants are present. The VOUS category corresponded to variants classified as “VOUS” at INFEVERS. Therefore, our dataset was represented by 41 PATHOGENIC, 89 VOUS, 41 BENIGN, and 45 UNRESOLVED variants. All variants were mapped along the *MEFV* coding sequence and its functional domains using Lollipops [14].

### Statistical comparison

We extracted the REVEL score of each variant and the mean, SD, and the 95% confidence intervals were determined for each category. The statistical significance of the differences between categories was evaluated by ordinary one-way ANOVA, and unpaired t-test between each category.

To evaluate the predictive potential of REVEL scores in the FMF context we built a classifier on the dataset represented by the BENIGN and PATHOGENIC variants (n=82). Five different algorithms were tested to identify the best model for predicting variants pathogenicity potential based on their REVEL score, namely, Linear Discriminant Analysis (LDA), Classification and Regression Trees (CART), k-Nearest Neighbors (kNN), Support Vector Machines (SVM) with a linear kernel, and Random Forest (RF). This set of algorithms was chosen as it represents a good combination of linear and non-linear models to test. The dataset was split into two parts, 80% was used to train the models through a 10-fold cross validation, and 20% was used as a validation dataset. All models were weighed using the “accuracy” metric. Computations were performed in R using the Caret package.

Finally, the specificity, sensitivity, accuracy, Matthews’ correlation coefficient and Youden index for each REVEL score value in correctly identifying BENIGN and PATHOGENIC variants were calculated. Where not otherwise stated GraphPad Prism V.8.0.1 (GraphPad Software, San Diego, California, USA) was used for all statistical analyses.

## Supporting information

**S1 Table. INFEVERS classification, REVEL scores, and proposed novel classification for 216 *MEFV* missense variants.** List of the 216 missense variants analyzed with previous and proposed classification.

**S1 File. Parameters used to assess the performances of different values of REVEL score.** The parameters used to evaluate the performance of different value of REVEL scores as threshold for pathogenicity are indicated with their results for the best performing threshold value of 0.298

